# Structure informed xrRNA2 mutations in Zika virus provide a roadmap for vaccine development

**DOI:** 10.64898/2026.05.27.728262

**Authors:** Camille Merrick, Imali Kegode, Sarah Leach, Madeline Kale, Dustin Heiden, J. David Beckham

## Abstract

Flaviviruses like Zika virus (ZIKV), contain RNA tertiary structures within the 3’ untranslated region (UTR) that halt the 5’-to-3’ RNA exonuclease, Xrn1. Halting of Xrn1 at the two RNA structures, termed exonuclease-resistantRNA1 and 2 (xrRNA1 and xrRNA2), results in the formation of subgenomic flavivirus RNAs (sfRNA) that support viral pathogenesis. While the role of the flavivirus xrRNA1 in pathogenesis is well characterized, the role of the flavivirus xrRNA2 structure is not well studied. Using xrRNA crystal structure data, we inserted structure-informed mutations in ZIKV xrRNA2 to disrupt tertiary folding independent of significant sequence changes, evaluate sfRNA production, and define pathogenesis in a murine model of ZIKV infection. Compared to our prior work with ZIKV xrRNA1, we found that ZIKV xrRNA2 is under increased selection pressure to maintain sfRNA production resulting in multiple targeted mutations in xrRNA2 junctional region to induce a stable mutant. Using three targeted xrRNA junctional mutations termed ZIKV X2.L1, we found that the resulting ZIKV clone exhibits attenuated cell death in cultures and decreased viral growth in tissue cultures. In a murine model of ZIKV infection, mice inoculated with ZIKV X2.L1 exhibit significantly decreased symptomatic infection, improved survival, decreased end-organ infection in the brain, and continued robust neutralizing antibody responses to ZIKV. Despite attenuation, serum from ZIKV X2.L1-infected mice or mice vaccinated with ZIKV X2.L1, exhibited 100% protection from lethal ZIKV challenge. These studies show that RNA structure-informed mutations provide a robust model for flavivirus attenuation and vaccine design. Additional studies defining the mechanisms of robust neutralizing antibody responses and flavivirus-specific vaccine development are needed to continue the development of this novel vaccine platform approach for medically important flavivirus infections.

**Author summary:** Zika virus is a member of the Orthoflavivirus (referred to as flavivirus) genus that exhibit conserved RNA structures in the 3’ untranslated region of the viral RNA genome. Two concerned RNA structures, termed exonuclease-resistant RNA 1 and 2, are important to support the ability of the virus to cause disease. While the first RNA structure is well studied, less is known about the role of exonuclease-resistant RNA 2 in the flavivirus infection. Using reverse genetics, we made mutations in the Zika virus exonuclease-resistant RNA 2 structure and studied how this mutant Zika virus was weakened or attenuated. We found that the mutant Zika virus clone exhibits reduced virus replication, reduced ability to kill cells, and decreased virulence in mouse models of Zika virus disease. Using this mutant virus as a potential vaccine candidate, we found that Zika virus with mutations in the exonuclease-resistant RNA 2 structure provide complete protection from lethal Zika virus challenge. These data suggest that targeting the second exonuclease resistant RNA structure in flaviviruses is a viable platform for the development of vaccine candidates for this important group of viruses.

## Introduction

Zika Virus (ZIKV) is a positive-sense, single-stranded, RNA virus belonging to the *Orthoflavivirus* (referred to as flaviviruses) genus in the *Flaviviridae* family. This genus includes other important human pathogens such as Dengue virus (DENV), West Nile virus (WNV), Japanese encephalitis virus (JEV), yellow fever virus, tick borne encephalitis virus, and many others (1). The flavivirus genome is characterized by a single open reading frame for structural and nonstructural proteins flanked by a 5’ and 3’ untranslated region (3’UTR) that play important functional roles in replication, translation, packaging and other roles in the viral life cycle (2). Instead of expressing a 3’ poly-A tail, the 3’UTR of flaviviruses exhibit several conserved RNA structures including duplicated stem-loop structures termed exoribonuclease - resistant RNA 1 (xrRNA1) and xrRNA2 at the 5’ end of the 3’ UTR (3). Xr-RNAs are defined by tertiary RNA interactions that result in specific three-dimensional (3D) structures that create a ring-like topology that is resistant to Xrn1-mediated RNA decay (4–6). The flavivirus xrRNA pseudoknot protects the remaining 3’UTR from host 5’ – 3’ exoribonuclease Xrn-1 RNA decay and maybe resistant to other host 5’ exoribonucleases as well (7). Once Xrn-1 halts at xrRNA1 or xrRNA2 the remaining 3’UTR functions as viral non-coding RNAs termed subgenomic flavivirus RNA 1 (sfRNA1) or sfRNA2, respectively. Mosquito-borne flaviviruses including WNV, Murray valley encephalitis virus, JEV, DENV, and ZIKV all exhibit xrRNA-dependent formation of sfRNAs that are required to support viral pathogenesis(3,4,6,8,9). Accumulation of sfRNAs during flavivirus infection supports viral replication, virulence, and pathogenesis(3,8,10–12). The kinetics and binding targets for sfRNAs result in epidemiologic fitness of DENV isolates (13). Thus, xrRNA-dependent sfRNA production is likely evolutionarily fine-tuned to the specific host interactions of individual flaviviruses.

The crystal structure of xrRNA1of ZIKV provided information on critical long-range nucleotide interactions required to stabilize the 3-D pseudoknot that were not previously predicted by secondary structural analysis(4). Using data from the crystal structure of ZIKV xrRNA, our laboratory previously demonstrated that a single point mutation to disrupt the phosphate-backbone interactions with the junctional loop of the xrRNA1 pseudoknot results in loss of sfRNA1 production and attenuated viral replication in the brain and placenta of a murine ZIKV infection model (14). While the role of ZIKV xrRNA1 in sfRNA production is well defined, the role of xrRNA2-dependent sfRNA biogenesis and viral pathogenesis not as well defined. Prior work has shown that ZIKV xrRNA2 interacts with Musashi-1 to support neuroinvasion (15). Another study demonstrated that xrRNA2 supports xrRNA1-dependent sfRNA production(16). While these studies provided important insights into the structure and function, structure-informed, clone-derived xrRNA2 mutations in rescued ZIKV have not been previously evaluated in viral pathogenesis studies.

Flavivirus-specific sfRNA function is dependent on the cell type specific interactions that are based on downstream cellular targets that differ between vertebrate and invertebrate cells. In this context sfRNAs function to inhibit mRNA decay, induce apoptosis, as well as, blunt type 1 interferon responses in mammalian cells and inhibit RNAi and toll pathways in invertebrate cells (2,13,17–19). During invertebrate cell infection, duplicated xrRNA structures in the 3’UTR of insect specific flaviviruses are redundant in nature to ensure that sfRNA is produced due to its importance during infection (20). In contrast, in human cells, xrRNA structures have been shown to have potential cooperation in the biogenesis of sfRNA species (21). These data suggest that structural and functional roles of sfRNAs and xrRNAs may be species specific during the host-switching life cycle of flaviviruses. However, specific studies evaluating xrRNA-specific function have not been rigorously evaluated.

Previous studies have shown that despite the diverse species tropisms between individual *Orthoflaviviruses*, xrRNAs have remarkably well conserved 3D structures in the 3’UTR (20,22). In this study we use the structural homology of xrRNA1 and xrRNA2 to make targeted mutations in the xrRNA2 structure to determine its role in ZIKV pathogenesis. We found that the single mutations targeting nucleotide interactions that stabilize the phosphate backbone kink in the xrRNA2 junction or targeting nucleotide interactions in the xrRNA loop that stabilize the 5’ end pseudoknot result in reversion. When these three mutations were combined, the resulting xrRNA2 mutant, termed ZIKV X2.L1, was found to be stable, exhibited loss of sfRNA2 production, and was able to replicate in cell culture. ZIKV X2.L1 exhibited significantly decreased cytopathic effect, increased sensitivity to interferon treatment, and was significantly attenuated in a murine model of ZIKV infection. Despite attenuation, ZIKV X2.L1 exhibited robust neutralizing antibody responses and serum from ZIKV X2.L1 infected mice protected naive mice from lethal ZIKV challenge. Lastly, we found that immunization with ZIKV X2.L1 provided 100% protection from lethal ZIKV challenge in type I interferon deficient mice. Taken together, these studies show for the first time that ZIKV xrRNA2-dependent sfRNA2 production is critically important for ZIKV virulence during mammalian infection and interferon escape resulting in attenuation with retained immunogenicity. The resulting live-attenuated ZIKV X2.L1 mutant provides robust protection as a vaccine candidate against lethal ZIKV challenge.

## Results

### Zika virus xrRNA1 and xrRNA2 exhibit different selection pressure

To determine the role of individual xrRNAs in ZIKV sfRNA biogenesis and pathogenesis, we first identified shared secondary RNA structures between xrRNA1 and xrRNA2 to predict overlapping function in the formation of Xrn1 resistance (**Fig. 1A**). Utilizing this approach, we identified a nucleotide mutation in xrRNA1 (C10415G) and xrRNA2 (C10496G) that disrupt the stabilizing interaction within the phosphate backbone kink of the loop structure (**Fig 1B**). Following isolation of respective ZIKV mutant viruses, RNA was isolated for 3’UTR sequencing and northern blot analysis using radioisotope-labeled probe targeted to the dumbbell region of the flavivirus 3’UTR. We found that ZIKV xrRNA1 C10415G mutation resulted in elimination of sfRNA1 production; however, the equivalent structural mutation in ZIKV xrRNA2 C10496G resulted in continued sfRNA1 and sfRNA2 biogenesis (**Fig 1C**). Sequencing of the respective mutant viruses showed that the C10415G xrRNA1 (X1) mutation was present but, the C10496G xrRNA2 (X2) mutation had reverted to the wildtype sequence. This data suggested that there is increased selection pressure on the ZIKV xrRNA2-dependent sfRNA2 biogenesis compared to xrRNA1 dependent sfRNA1 biogenesis.

**Fig. 1.**
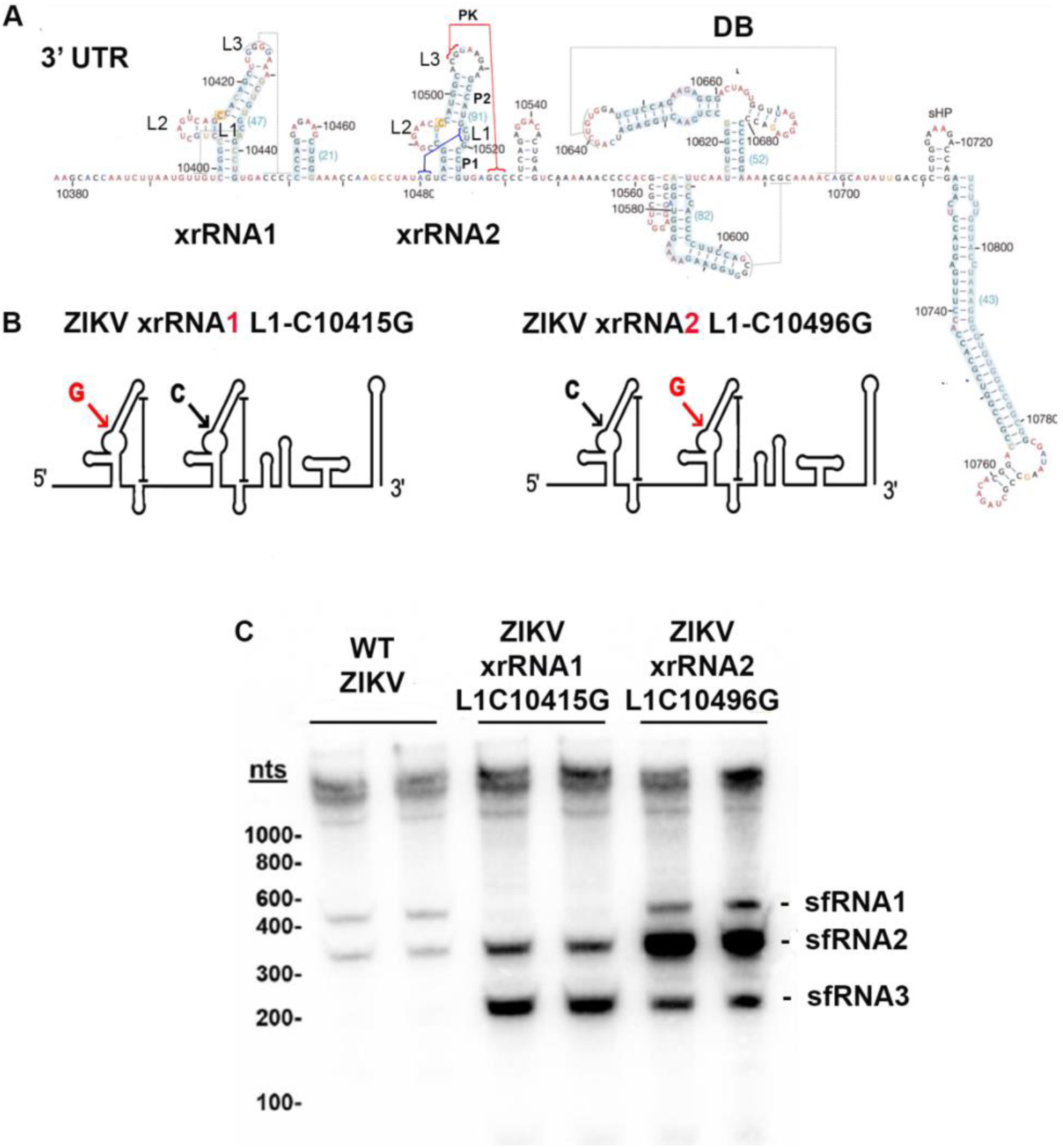
Zika virus xrRNA2 exhibits increased resistance to mutagenesis compared to xrRNA1. (**A**) Diagram of ZIKV 3’ UTR showing crystal-structure derived predictions of xrRNA2 L1 interactions with the 5’ end of the structure (Blue). (**B**) Diagram of mutation locations in ZIKV xrRNA1 (C104415G) and ZIKV xrRNA2 (C10496G) predicted to disrupt tertiary structure interaction with the phosphate backbone of the pseudoknot and allow for Xrn1 degradation. (**C**) Representative northern blot (n=2) of RNA from Vero cells following infection with WT ZIKV, ZIKV xrRNA1 (C10415G) in L1, and ZIKV xrRNA2 (C10496G) in L1 (MOI 1, 48 hours post-infection). 3’UTR radiolabeled to detect sfRNA production.

### Disruption of ZIKV L1 to 5’ xrRNA2 interactions results in a small plaque phenotype

To further define the selection pressure to maintain the ZIKV xrRNA2 pseudoknot, point mutations C10518G and U10519A we introduced into the L1 junctional loop of xrRNA2 to disrupt L1 interactions with the 5’ end of xrRNA (**Fig 2A-C**). Following transfection and virus isolation, RNA was isolated for 3’UTR sequencing showing that only the U10519A mutation (L1 U,A) was stable (**Fig 2C**). To determine if ZIKV L1U,A exhibited phenotypic differences compared to WT-ZIKV, A549 cells were infected at a multiplicity of infection (MOI) 0.1 with WT ZIKV or ZIKV L1U,A, and cell lysate and supernatant were collected at 0, 24, 48 and 72 post infection. We found no significant difference between WT-ZIKV and ZIKV L1U,A in virus replication as measured by genome copies or infectious viral titer in supernatant or cell lysates (**Fig 2D-F**). Despite the similar growth kinetics between WT-ZIKV and ZIKV L1U,A, we observed a small plaque phenotype in ZIKV L1U,A (-3.5fold plaque size) compared to WT-ZIKV (n=6, p=0.0004, two-way ANOVA, **Fig. 2G,H**). These data suggest that a single nucleotide change in the xrRNA2 loop is sufficient to result in viral phenotypic changes but additional stabilizing mutations in the xrRNA2 loop are likely required to support stable loss of xrRNA2 function.

**Fig. 2.**
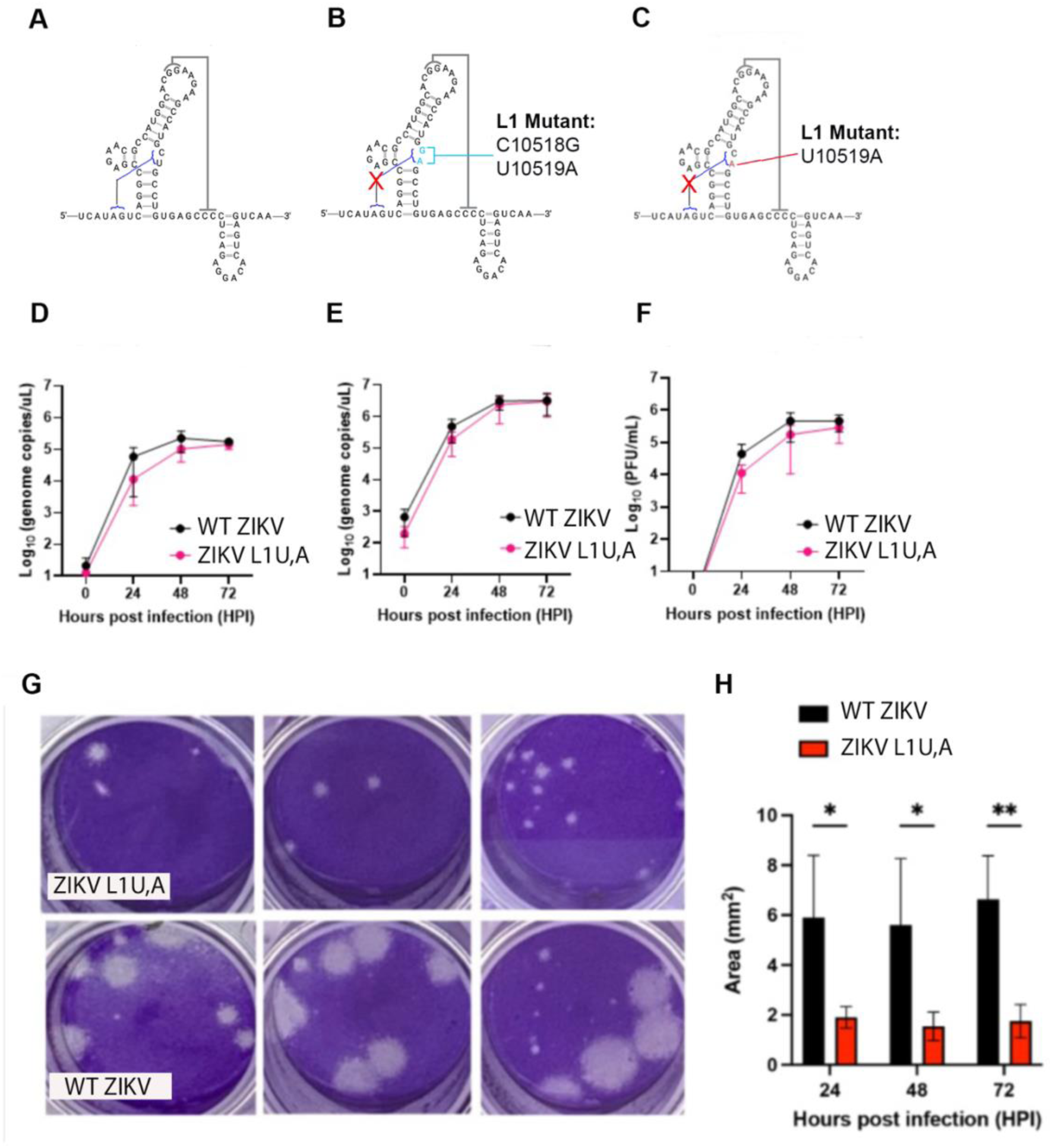
Disruption of Zika virus xrRNA2 L1 (U10510A) with the 5’ end results in a small plaque phenotype. We produced clone-derived ZIKV virus isolates with (**A**) WT ZIKV xrRNA2, (**B**) ZIKV xrRNA2 mutations at C10518G, U10510A, and (**C**) ZIKV xrRNA2 U10519A. ZIKV xrRNA2 (C10518G, U10510A) reverted to ZIKV xrRNA2 U10510A. Vero cells were inoculated with WT ZIKV or ZIKV xrRNA2 U10510A (referred to as ZIKV L1U,A) at a MOI of 1.0. There was no significant difference in ZIKV genome copies from (**D**) supernatant or (**E**) cell lysates comparing WT ZIKV and ZIKV L1U,A. (**F**) Infectious plaque forming units (PFU) exhibited no significant difference in supernatant from WT ZIKV and ZIKV L1U,A inoculated vero cells. (**G,H**) ZIKV plaques from ZIKV L1U,A exhibited significantly smaller plaque size compared to WT ZIKV plaques. *p<0.05, **p<0.01, Student t-test.

### Targeted nucleotide mutations in ZIKV xrRNA2 eliminate sfRNA2 production

Next, we introduced ZIKV xrRNA2 mutations that disrupt the L1 phosphate backbone kink (C10496G) and distal L1 (C10518G and U10519A) interactions with the 5’ end of the xrRNA2 structure, termed X2.L1 ZIKV (**Fig. 3A-B**). Following transfection and isolation of virus, we found that X2.L1 ZIKV exhibited a 125-fold decrease in supernatant genome copies compared to WT-ZIKV (n=3, p<0.0001, two-tailed T-test, **Fig 3C**). Following passage of X2.L1 ZIKV in C6/36 cells, viral titer was determined using FFU. We found that X2.L1 ZIKV exhibited a 4.35-fold decrease in FFU formation compared to WT-ZIKV (n=6, p=0.0002, t-test, **Fig 3D**). Sequencing of X2.L1 ZIKV stock revealed stable mutations and no evidence of compensatory mutations in the xrRNA2 structure. Next, we determined the role of X2.L1 ZIKV on sfRNA2 biogenesis using an in vitro Xrn1 digest of the X2.L1 ZIKV 3’UTR vs WT ZIKV 3’UTR, both of which lacked expression of xrRNA1 at the 5’ end so xrRNA2 function could be evaluated in isolation of xrRNA1. Following RNA folding, we found that the 3’UTR of X2.L1 ZIKV was unable to halt Xrn1 and protect the 3’UTR while WT-ZIKV xrRNA2 3’UTR maintained sfRNA biogenesis (n=2, **Fig. 3E**).

**Fig. 3.**
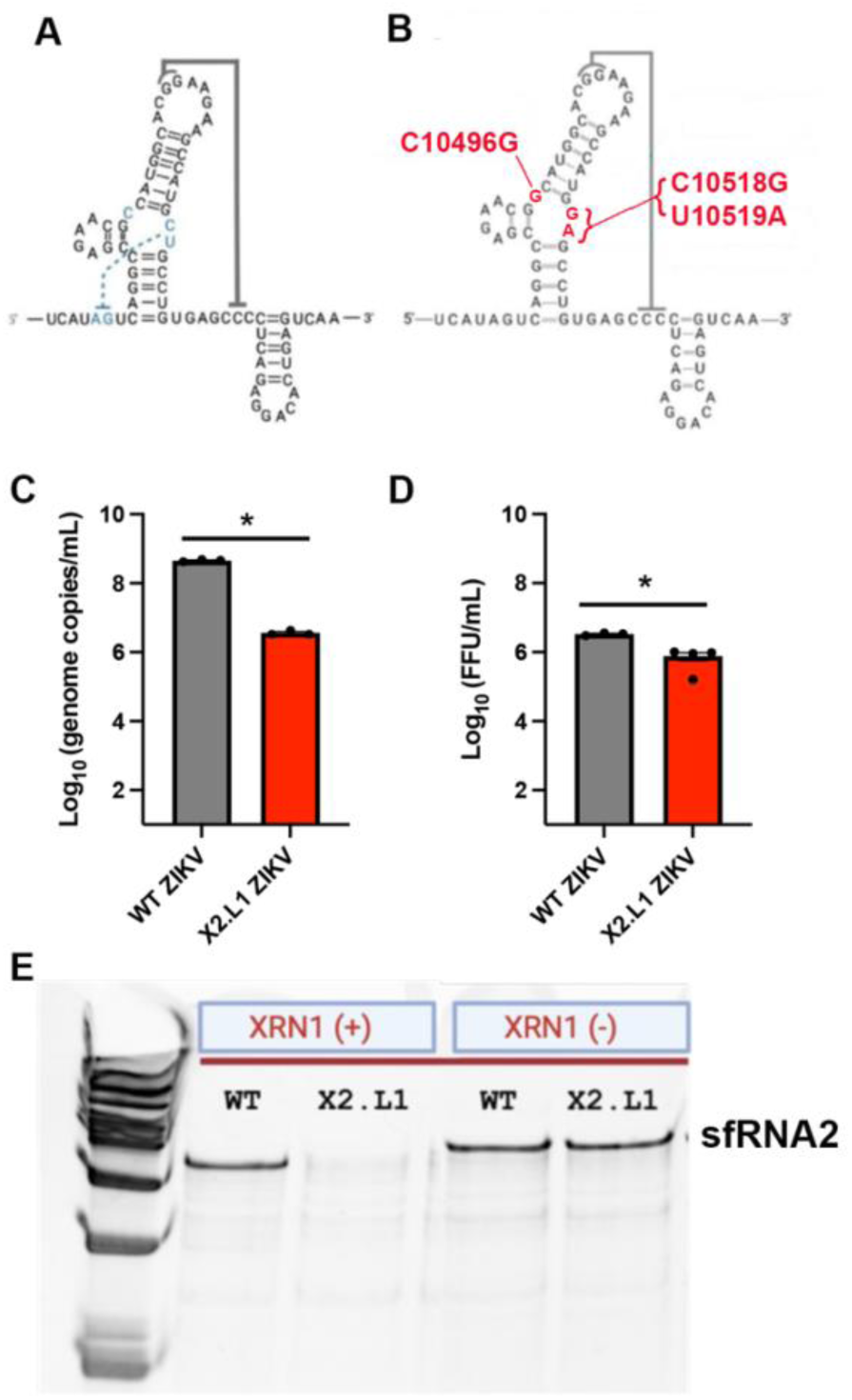
Targeted nucleotide mutations in ZIKV xrRNA2 eliminate sfRNA2 production. We produced clone-derived ZIKV virus isolates with (**A**) WT ZIKV xrRNA2, (**B**) ZIKV xrRNA2 mutations at C10496G, C10518G, and U10510A, termed X2.L1 ZIKV. (**C,D**) X2.L1 ZIKV exhibited stable sequencing with no reversion, ZIKV genome copy number, and ZIKV infectious particles that were slightly decreased compared to WT ZIKV. *p<0.05, t-test. (**E**) Gel image of a representative *in vitro* Xrn1 RNA degradation assay of WT ZIKV 3’UTR compared to X2.L1 ZIKV 3’UTR showing loss of sfRNA2 with X2.L1 mutations in the xrRNA2 structure (n=2).

### X2.L1 ZIKV is attenuated in cell culture

We next determined the replication kinetics, cell injury, and interferon sensitivity of X2.L1 ZIKV. A549 cells were inoculated with WT ZIKV or X2.L1 ZIKV (MOI 0.1) and cell supernatant and cell lysate were collected at 0, 24, 48 and 72 hours post infection. We found a significant decrease in genome copies and infectious viral titer of X2.L1 ZIKV at 48 hours post infection compared to WT-ZIKV (n=6, p=0.001, two-way ANOVA, **Fig. 4A-C**) and all other time points were similar. The plaque phenotype of X2.L1 ZIKV resulted in poorly defined plaques, so FFU was utilized for infectious viral titer assays. To test attenuation in vitro, A549 cells were inoculated with WT ZIKV or X2.L1 ZIKV (MOI=1.0) and cell injury was measured using an XTT assay at 24, 48 and 72 hours post infection. X2.L1 Relative to mock-inoculated cells, ZIKV inoculated cells exhibited a significant decrease in cellular injury at 48 (p<0.0001) and 72 hours post-infection compared to WT ZIKV inoculated cells (n=6, p=0.001, two-way ANVOA, **Fig. 4D**). Additionally, X2.L1 ZIKV inoculated cells exhibited no significant injury beyond mock-inoculated cells. These data are similar to prior work showing that ZIKV xrRNA-dependent sfRNA production is critical to support virus-induced cell injury (14,23). Prior studies have shown that alterations to flavivirus sfRNA1 production result in a reduced ability to escape interferon restriction (3,9,24,25). To determine if the X2.L1 ZIKV exhibited a change in type 1 interferon sensitivity due to reduced sfRNA production, vero cells were inoculated with WT ZIKV or X2.L1 ZIKV (MOI=1.0) and treated with serial dilutions of interferon-alpha2, followed by harvest of cells for genome copies at 48hours post-infection. The total genome copies were determined for each dilution of interferon-alspha2 and an inhibitory concentration (IC) that inhibited 50% of virus genome replication (IC50) was calculated for each replicate. We found that WT ZIKV (IC50=91.44) exhibited increased resistance to interferon-alpha2 compared to X2.L1 ZIKV (IC50=30.9, n=10, Least squares regression). Together, these data show that X2.L1 ZIKV exhibits decreased viral titer, decreased cell injury, and increased sensitivity to type I interferon compared to WT ZIKV.

**Fig. 4.**
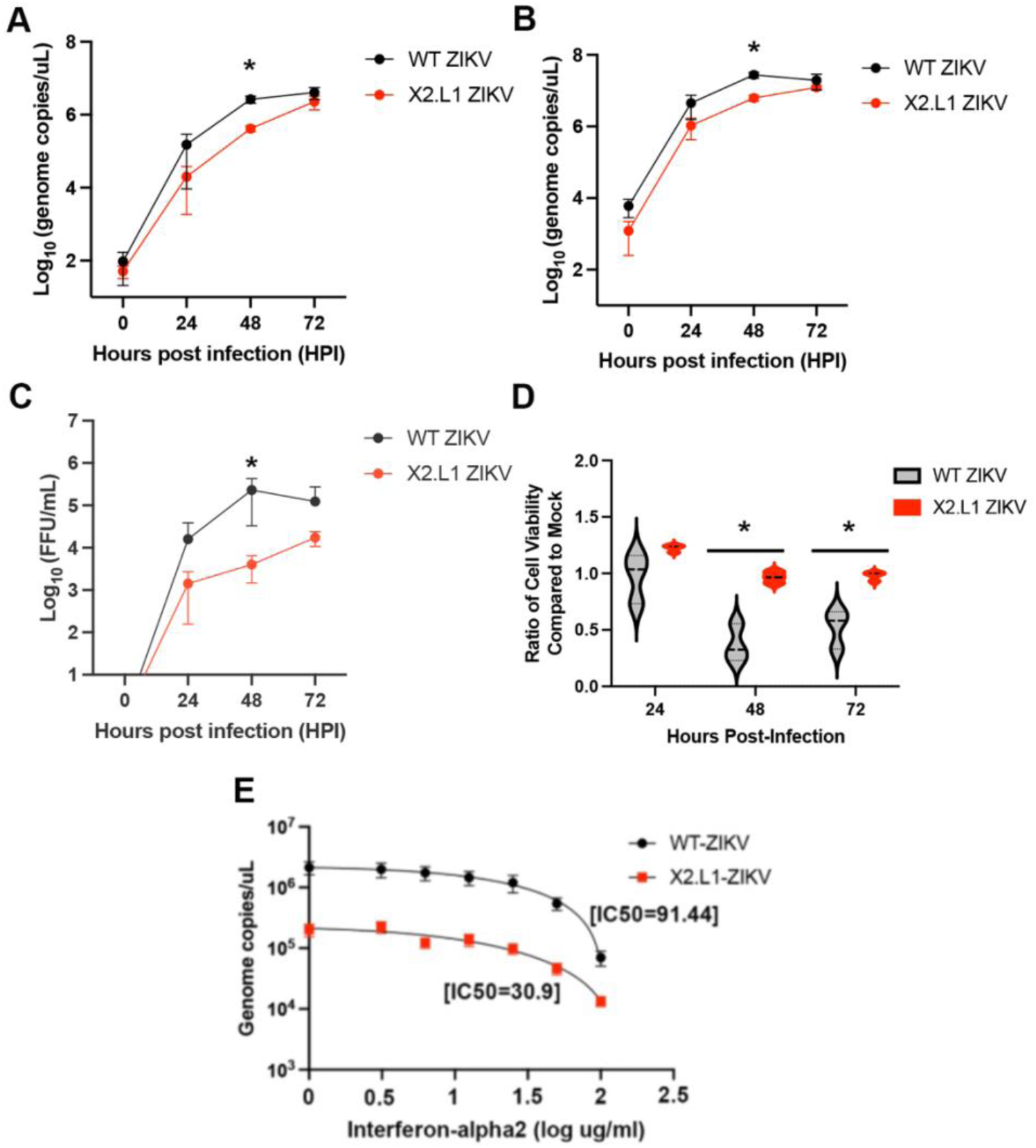
X2.L1 ZIKV exhibits attenuation in cell culture. Vero cells were inoculated with WT ZIKV or X2.L1 ZIKV at a MOI of 0.1 followed by analysis at indicated time points for (**A**) ZIKV genome copies in the supernatant, (**B**) ZIKV genome copies in cell lysates, and (**C**) infectious ZIKV particles in the supernatant as measured by focus forming units (FFU). *p=0.001 two-way ANOVA. (**D**) Vero cells were inoculated with WT ZIKV or X2.L1 ZIKV (MOI=1) and cell injury was measured at the indicated time points using an XTT assay. *p<0.001, two-way ANOVA. (**E**) Vero cells were inoculated with WT ZIKV or X2.L1 ZIKV (MOI=1) and treated with serial dilutions of interferon-alpha2 followed by determination of genome copies at 48 hours post-infection. Using the dilution curve of interferon-alpha2, we determined the concentration of interferon required to inhibit 50% ZIKV genomes (IC50). X2.L1 ZIKV exhibited an IC50=30.9 and WT ZIKV exhibited an IC50=91.44. N=10, Least squares regression.

### X2.L1 Zika virus is attenuated in a mouse model of disease

Next, we determined if X2.L1 ZIKV was attenuated in immune deficient murine models of ZIKV infection. First, we inoculated type I interferon receptor knockout (IFNAR1 KO) mice with either WT ZIKV or X2.L1 ZIKV (10,000 PFU, intraperitoneal (ip) inoculation) and mice were euthanized at 8 days post-infection for analysis of ZIKV genome copies in the brain and spleen tissue. In IFNAR1 KO mice, WT and X2.L1 ZIKV inoculated mice exhibited no significant weight loss compared to mock-inoculated control IFNAR1 KO mice (**Fig. 5A**). We also found no significant difference in WT ZIKV and X2.L1 ZIKV genome replication in brain or spleen tissue of IFNAR1 KO mice (Brain: n=6, p=0.0537(n.s.), one-way ANOVA, **Fig. 5B**; spleen: n=6, p>0.05(n.s.), one-way ANOVA, **Fig. 5C**). We next defined the difference in WT and X2.L1 ZIKV (10,000 PFU, ip) pathogenesis in type I and type II interferon receptor knockout (AG129) mice. We found that AG129 mice inoculated with WT ZIKV exhibited significantly increased (10-20%) weight loss compared to mock and X2.L1 ZIKV inoculated mice (*p<0.0001, n=18, Two-way ANOVA, repeated measures). At day 8 post-infection, brain and spleen tissue was collected from these mice, and RNA was extracted to run qPCR to assess viral genome copies. We found no significant difference in viral genome copies between WT-ZIKV and X2.L1 ZIKV in the spleen (n=6, one-way ANOVA with multiple comparisons, **Fig. 5E**). However, we found a 90% decrease in viral genome copies in the brains of AG129 mice inoculated with X2.L1 ZIKV (mean 94,643 ZIKV genome copies/mg tissue) compared to WT-ZIKV (mean 963,478 ZIKV genome copies/mg tissue; n=6, p <0.0001, one-way ANVOA with multiple comparisons, **Fig. 5F**). As the clone derived wildtype ZIKV derived from PRVABC59 did not exhibit a significant phenotype in IFNAR KO mice, we next evaluated X2.L1 compared to a strain with virulence in murine models: Zika Paraiba 01 Brazil 2015 (WT-ZIKV Brazil). IFNAR1-KO mice were inoculated with 10,000 PFU ip with WT-ZIKV Brazil and X2.L1 ZIKV and followed for 21 days. We found that IFNAR KO mice inoculated with WT-ZIKV Brazil exhibited significantly increased weight loss following infection compared to IFNAR KO mice inoculated with X2.L1 ZIKV (p=0.0014, Two-way ANOVA, repeated measures, **Fig. 5G**). Following inoculation, IFNAR KO mice were evaluated for survival as defined by development of infection morbidity due to encephalitis requiring euthanasia or weight loss >20%. We found that mice inoculated with WT ZIKV Brazil exhibited significantly decreased survival (median survival 12.5 days) compared to X2.L1 ZIKV inoculated mice (median survival undefined, p=0.0179, Logrank test, **Fig. 5H**). Compared to a virulent ZIKV isolate, we found that X2.L1 ZIKV exhibits robust attenuation in a murine model of ZIKV infection.

**Fig. 5.**
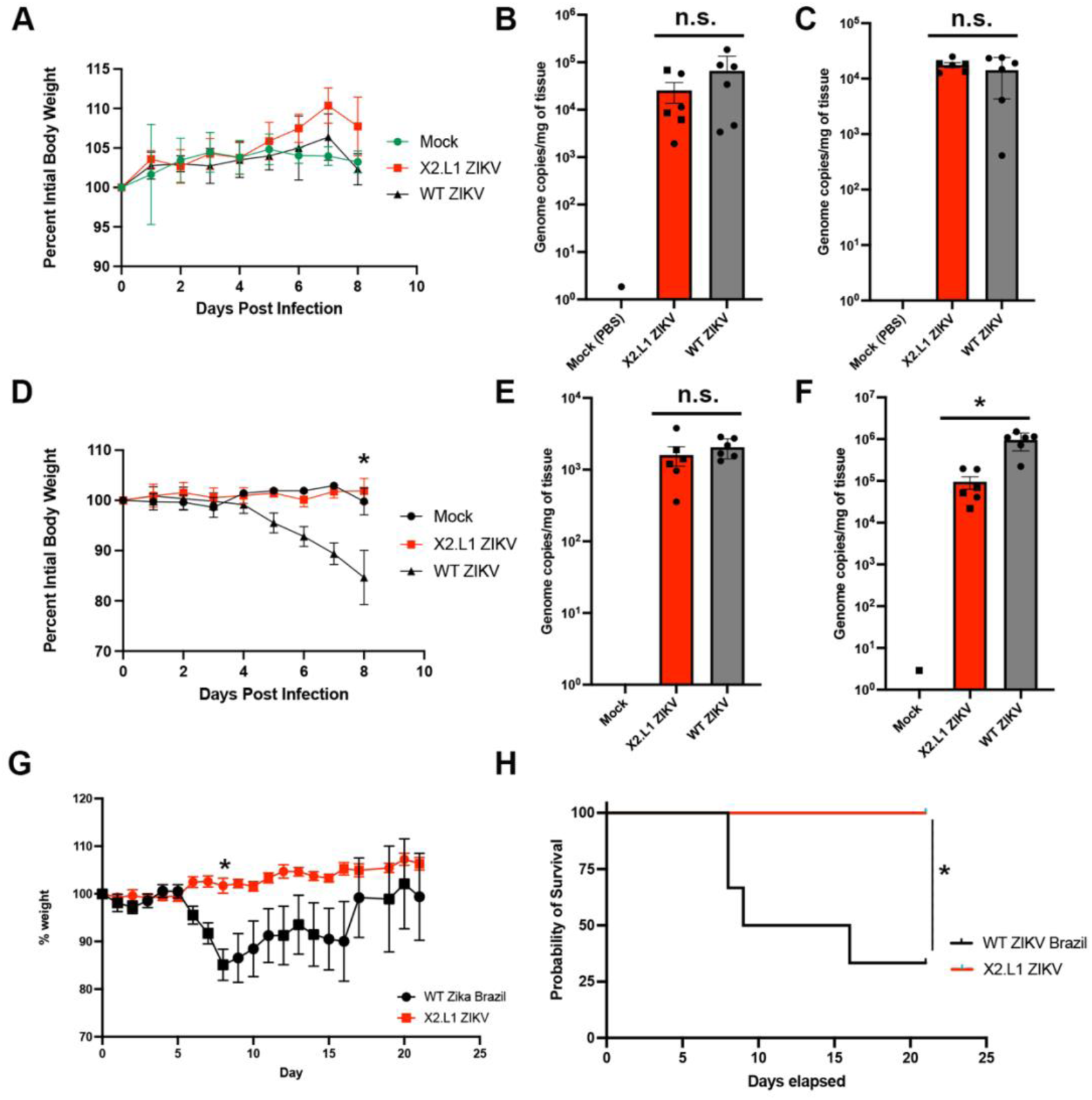
X2.L1 ZIKV is Attenuated in Immune Compromised Mice Type I interferon receptor knockout (IFNAR1 KO) mice were inoculated with mock control, WT ZIKV, or X2.L1 ZIKV (10^4^ pfu, intraperitoneal [ip] inoculation), and mice were euthanized at day 8 post-infection. (**A**) There was no significant difference in weight changes between mock, WT ZIKV, and X2.L1 ZIKV inoculated mice. At day 8 post-infection, there was no significant difference in ZIKV genome copy number between mock, WT ZIKV, and X2.L1 ZIKV in the (**B**) brain or (**C**) spleen tissue (n=6, *p>0.05(n.s.), one-way ANOVA). Next, we inoculated type I and type II interferon receptor knockout (AG129) mice with mock control, WT ZIKV, or X2.L1 ZIKV (10^4^ pfu, intraperitoneal [ip] inoculation), and mice were euthanized at day 8 post-infection. (**D**) AG129 mice inoculated with WT-ZIKV exhibited 10-20% increased weight loss compared to mock and X2.L1 ZIKV inoculated mice (*p<0.0001, n=18, two-way ANOVA, repeated measures). At day 8 post-infection, (**E**) there was no difference in ZIKV genome copy number in the spleens of WT ZIKV and X2.L1 ZIKV inoculated AG129 mice; however, (**F**) AG129 mice inoculated with X2.L1 ZIKV exhibited a 90% decrease in mean ZIKV genome copies in brain tissue compared to WT ZIKV inoculated mice (n=6, *p<0.0001, one-way ANOVA with multiple comparisons). Next, we inoculated IFNAR KO mice with Zika Paraiba 01 Brazil 2015 (WT ZIKV Bazil) compared to X2.L1 ZIKV (10^4^ pfu, ip inoculation). We found that WT ZIKV Brazil inoculated mice exhibited significantly increased (**G**) weight loss (n=6, *p=0.0014, two-way ANOVA, repeated measures) compared to X2.L1 inoculated mice. (H) Additionally, X2.L1 ZIKV inoculated mice exhibited significantly decreased mortality (*p=0.0179, Logrank test) compared to WT ZIKV Brazil inoculated IFNAR KO mice.

### X2.L1 ZIKV Inoculation Produces Protective Neutralizing Antibody Responses

Since X2.L1 ZIKV exhibited attenuation in tissue culture and in murine models compared to WT-ZIKV, we next determined if X2.L1 ZIKV exhibited protective neutralizing antibody responses despite attenuation. First, IFNAR1-KO mice were inoculated with 10,000 PFU IP of WT ZIKV, X2.L1 ZIKV, or PBS and serum was collected at 21 days. Total anti-ZIKV antibody in serum samples revealed an increase in ZIKV-specific antibody in WT ZIKV and X2.L1 ZIKV inoculated mice (One-Way Anova, p<0.0001, **Fig. 6A**). Since absolute ZIKV-specific antibody responses following X2.L1 ZIKV inoculation were not statistically different from mock inoculated mice, we next determined ZIKV neutralization capacity of serum from inoculated mice. Using serial dilutions of serum from WT ZIKV and X2.L1 ZIKV inoculated mice, we found that WT ZIKV (FRNT50=1:180,000) and X2.L1 ZIKV (FRNT50=1:6,024) exhibit robust neutralization compared to mock-inoculated mice (**Fig. 6B**). Next serum from X2.L1 ZIKV inoculated IFNAR1-KO mice and non-infected IFNAR1-KO mice was pooled and AG129 mice were inoculated with 100uL of serum (ip). 24 hours later mice were inoculated with 10,000 PFU of WT-ZIKV Brazil subcutaneously (SC) and mice were monitored for symptoms and weight loss over 15 days (**Fig. 6C**). On day 7 post-infection, all AG129 mice treated serum from non-infected mice had lost >20% starting weight and develop moribund symptoms of ZIKV encephalitis requiring euthanasia (median survival 8 days); instead, AG129 mice treated with serum from X2.L1 ZIKV mice exhibited minimal weight loss and 100% survival (p=0.0009, Logrank test, **Fig. 6D,E**).

**Fig. 6.**
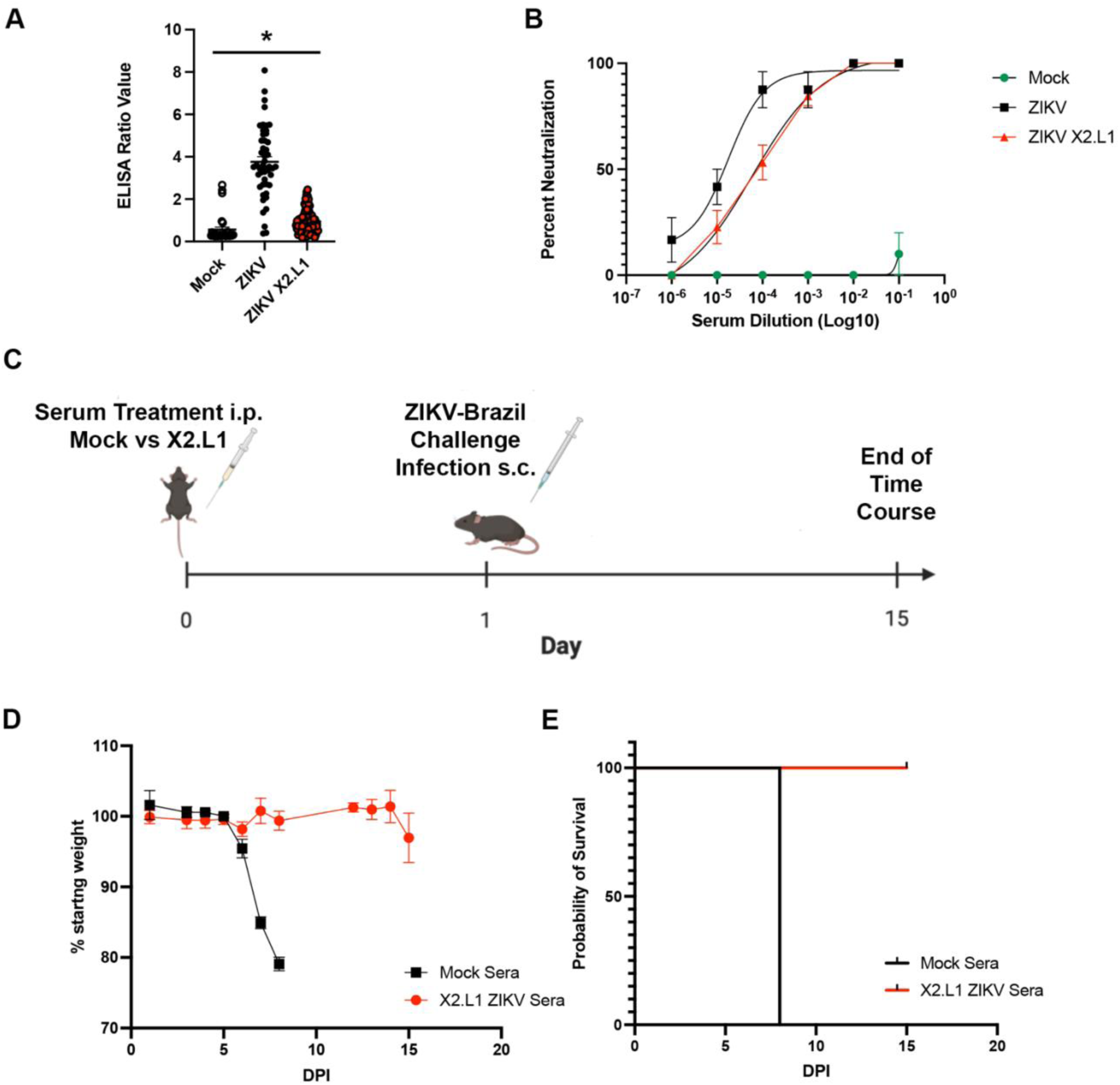
X2.L1 ZIKV Inoculation produces protective neutralizing antibodies IFNAR KO mice were inoculated with mock, WT ZIKV, or X2.L1 ZIKV (10^4^ pfu, ip) and euthanized at day 21 post-infection for serum collection. (**A**) Total anti-ZIKV antibody were significantly increased (*p<0.0001, one-way ANOVA, multiple comparisons) in WT ZIKV inoculated mice and there was a non-statistically significant increase in X2.L1 ZIKV inoculated mice compared to mock-inoculated mice. (**B**) Neutralization antibody titers were determined in the same experimental groups using focus-forming unit neutralization 50 (FRNT50) assays showing that both WT-ZIKV (FRNT=1:180,000) and X2.L1 ZIKV (FRNT=1:6,024) exhibited high titer neutralization in the serum. (**C**) Next, serum from mock inoculated and X2.L1 inoculated mice was transferred to AG129 mice (100microL of serum ip) and mice were inoculated with WT ZIKV Brazil after 24 hours. (**D**) AG129 mice inoculated with X2.L1 ZIKV serum exhibited protection from weight loss following lethal WT ZIKV Brazil challenge (10^4^ pfu, subcutaneous inoculation) compared to AG129 mice inoculated with serum from infected serum and (**E**) X2.L1 ZIKV serum protected naïve AG129 mice from WT ZIKV Brazil mortality (n=6, *p=0.0009, Logrank test).

### Vaccination with ZIKV X2.L1 Protects Immune Deficient mice from ZIKV Challenge

To determine the role of X2.L1 ZIKV vaccination in the protection of mice from lethal ZIKV challenge, IFNAR1-KO mice were inoculated with mock or ZIKV X2.L1 (10,000 PFU ip) and after 21 day challenged with WT-ZIKV Brazil (10,000 PFU, SC, **Fig. 7A**). We found that IFNAR1 KO mice vaccinated with PBS exhibited significantly increase weight loss compared to ZIKV X2.L1 inoculated IFNAR1 KO mice following challenge with WT ZIKV Brazil (n=24, p<0.0001, Two-way ANOVA with multiple comparisons, **Fig. 7B**). Additionally, we found that IFNAR1 KO mice inoculated with PBS control exhibited a median survival of 11.5 days compared to 0% mortality in IFNAR1 KO mice inoculated with ZIKV X2.L1 (Logrank test, p=0.0018, **Fig. 7C**). To determine the degree of vaccine protection from ZIKV replication in the ZIKV X2.L1 vaccination model, we next collected brain and spleen tissue from mice 8 days after challenge with WT-ZIKV Brazil and RNA was extracted for analysis of ZIKV genome copies. We found that IFNAR1 KO mice vaccinated with X2.L1 ZIKV exhibited significantly decreased (mean=133 copies/mg tissue) ZIKV genome copies in spleen tissue compared to PBS-vaccinated mice (mean=786 genome copies/mg tissue, n=6, p=0.0234, **Fig. 7D**). Similarly, brain tissue from ZIKV X2.L1 inoculated mice exhibited no detectable virus compared to PBS vaccinated mice that exhibited a mean of 87,447 genome copies/mg of tissue (n=6, p=0.0373, **Fig. 7E**). These data show that ZIKV X2.L1 was completely protected immune compromised mice from lethal infection by preventing infection of brain tissue.

**Fig. 7.**
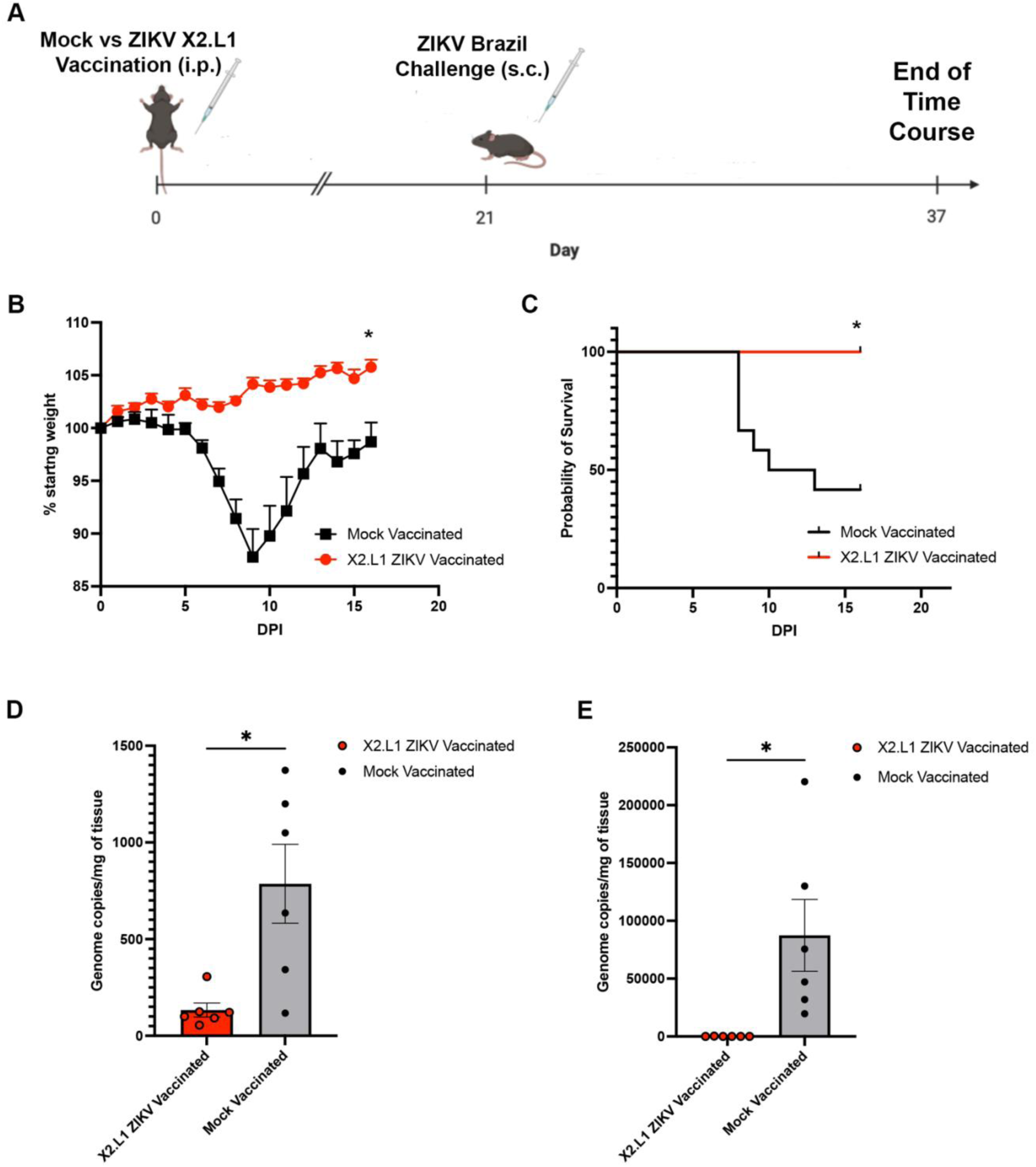
Vaccination with X2.L1 ZIKV protect Immunocompromised mice from ZIKV challenge. (**A**) IFNAR1 KO mice were inoculated with mock or X2.L1 ZIKV (10^4^ pfu, ip) followed by WT ZIKV Brazil (10^4^ pfu, sc) challenge at day 21 post-vaccination. (**B**) X2.L1 ZIKV vaccinated mice exhibited no significant weight loss following WT ZIKV Brazil challenge compared to mock vaccinated animals (n=12, *p<0.0001, two-way ANOVA with multiple comparisons). (**C**) IFNAR KO mice inoculated with mock vaccination exhibited a median survival of 11.5 days compared to 0% mortality in X2.L1 ZIKV vaccinated mice following WT ZIKV Brazil challenge (n=12, *p=0.0018, Logrank test). At 8 days post-infection, mice vaccinated with mock or X2.L1 ZIKV and challenged with WT ZIKV Brazil were euthanized for ZIKV genome copy analysis in the spleen and brain tissue. (**D**) X2.L1 ZIKV vaccinated mice exhibited significantly decreased ZIKV genome copies in the spleen (n=6, *p=0.0234, t-test) and (**E**) in brain tissue (n=6, *p=0.037, t-test).

## Discussion

In *Orthoflaviviruses*, xrRNA1 and xrRNA2 at the 5’ end of the 3’UTR stall the host 5’-to-3’ exonuclease Xrn1 leading to the formation of sfRNA1 and sfRNA2 (3). The role of xrRNA duplications is not well defined. The xrRNA pseudoknots may contribute equally to the formation of sfRNA1 and sfRNA2 to maintain sfRNA production through redundancy; however, recent studies have also suggested that there are inter-structural interactions that result in virus-host species specific sfRNA outcomes to support the unique pathogenesis of the individual *Orthoflaviviruses* (20,21). Our data show that ZIKV xrRNA2 exhibits a higher barrier to disruption compared to xrRNA1. We have previously show that a single mutation C-to-G mutation at position 22 in xrRNA1 was maintained through multiple passages, eliminated production of sfRNA1, and was maintained during infection in a murine model of ZIKV disease(14). However, the analogous mutation in ZIKV xrRNA2 (C10496G) in this study resulted in reversion of the mutation and continued sfRNA2 production during virus isolation. To maintain xrRNA2 disruption, we added additional xrRNA2 mutations in the distal L1 loop (C10518G and U10519A) to disrupt interactions with the 5’ end of the xrRNA2 structure, termed X2.L1 ZIKV. This strategy resulted in loss of sfRNA2 production and a decrease in infectious virus titer despite unchanged genome replication. Thus, targeted disruption of the xrRNA2 structure introduced in this study exhibited evidence of reduced viral titer independent of viral genome replication. Based on these data, we concluded that disruption of xrRNA2-dependent sfRNA2 production resulted in a decrease in infectious virions with no change in viral genome replication. Since mutations of ZIKV xrRNA2 exhibited an altered plaque phenotype, we expect that decreased infectious titer was secondary to decreased virulence following infection and not secondary to defects in genome replication.

Other studies that have evaluated *Orthoflavivirus* xrRNA2 have utilized deletions of the entire structure to determine function (3,21). Deletion xrRNA structures may not identify the structural roles of the second stem loop structure since the duplication may be able to maintain the function of a single or terminal xrRNA structure interactions with other 3’UTR structures like the dumbbell regions. Due to structural conservation between xrRNA1 and xrRNA2, the X2.L1 mutation introduced in the study should significantly disrupt the tertiary folding of xrRNA2 creating significantly increased distance between xrRNA1 and the downstream dumbbell structure or may alter the function of sfRNA1 since the xrRNA2 structure is included in the first sfRNA. Thus, the duplication of xrRNAs and specifically the most 3’ xrRNA may play an important role in mechanical protection of the downstream 3’UTR structures including xrRNA2 to maintain functional cellular interactions with sfRNAs that support distinct interactions with sfRNAs based on the expression of xrRNA in the sfRNA. More studies are needed to understand the functional role of xrRNAs included in sfRNAs to determine any interactions that are leading to the increase selection pressure on the terminal xrRNA structure.

Most mosquito-borne *Orthoflaviviruses* have at least 2 xrRNA structures resulting in the production of at least 2 sfRNA species (7,21). Studies investigating the functional differences between the different length sfRNAs in dengue have suggested that sfRNAs play an important role in host switching between invertebrates and mammalian hosts (8). For Zika Virus, we have previously shown that a mutation in xrRNA1 leading to subsequent loss of sfRNA1 was tolerated in both mammalian, A549, and mosquito, U4.4, cells (14). Despite expressing an isolated sfRNA 2, we have shown that disruption of xrRNA1-dependent sfRNA production results in moderate loss of virulence in cell cultures and in a murine model of ZIKV disease. Interestingly, yellow fever virus and Dengue 4, express a single xrRNA structure and produce an effective sfRNA species (26,27). Thus, the duplication of the xrRNA structures may be a mechanism to ensure expression of a terminal xrRNA structure so that biogenesis of sfRNAs can include at least one xrRNA.

Since X2.L1 ZIKV exhibited decreased virulence, we next evaluated the mechanism. Once sfRNA biogenesis begins, it is known to modulate host antiviral mechanisms including type 1 interferon response in vertebrates, RNAi and toll pathways in insects, cellular mRNA decay, viral RNA sensing, and interferon stimulated genes (9,13,19,28–32). Our studies show that X2.L1 ZIKV exhibited increased sensitivity to interferon treatment and was markedly reduced in virulence in tissue culture. We have previously shown that ZIKV dumbbell mutations result in decreased overall expression of sfRNAs following infection which resulted in marked decrease in virulence of ZIKV dumbbell mutants as measured by cell death assays and activated caspase-3 assays (23). Dengue virus-2 sfRNA induces Bcl-2 associated apoptosis in mammalian cells (12). However, ZIKV sfRNA inhibits apoptosis in infected mosquitoes and loss of sfRNA production results in activation of caspase 7 (17). These results indicate that sfRNA production modulates cell death and injury in a host-specific manner. In our studies, we find that ZIKV sfRNA production increases cell death and this supports viral pathogenesis in mammalian tissue culture and in murine models. Following inoculation of ZIKV or related flaviviruses in a mammalian host, initial replication occurs in local keratinocytes, Langerhans’s cells, dendritic cells, and monocytes which are required for dissemination (33). Dendritic cells and related CD45+ leukocytes are also important antigen presenting cells for the adaptive immune response, and ZIKV-induced apoptosis of these cells during viral pathogenesis may play an important role in the inhibition of the adaptive immune response during mammalian infection. Further studies are needed to evaluate the cell-type specific role of sfRNA expression during flavivirus pathogenesis.

Our studies provide new insight into the role of xrRNA-dependent sfRNA expression in neuroinvasion and neurovirulence in the central nervous system (CNS). We found that X2.L1 ZIKV exhibited similar replication in the brain and spleen in type 1 interferon receptor knockout mice. However, in mice with type I and type II interferon receptor deletion (AG129), we found that X2.L1 exhibited a 90% reduction in viral genome expression in the CNS while exhibiting unchanged replication in the spleen compared to WT ZIKV. These data show that xrRNA2-dependent sfRNA production is important for viral infection and replication in the CNS in the absence of type I and type II interferon restriction. This suggests that flavivirus sfRNA expression may play a role in escape of type III interferon restriction at the blood-brain-barrier (BBB). While there has been some research on what kinds of cells ZIKV is infecting to cross the placenta, the mechanism it uses to cross the BBB is unclear. The type III IFN response has been shown to be an effective antiviral response in barrier cells (28). The endothelium cells of the BBB utilize type 3 IFN to improve barrier function as opposed to systemic inflammatory responses associated with type 1 IFN (34). More studies will be needed to define the role of flavivirus sfRNA expression in the pathogenesis of BBB function and type III interferon inhibition.

Despite the attenuation of X2.L1 ZIKV in cell culture and in murine models of ZIKV infection, mice inoculated with X2.L1 developed neutralizing antibody activity that was similar to WT ZIKV infection. Additionally, we found that serum from X2.L1 ZIKV inoculated mice provided complete protection from lethal ZIKV-Brazil challenge in AG129 mice. Taken together, these data show that neutralizing antibodies to ZIKV provide an immune correlate of protection against lethal ZIKV infection in our studies. Our findings are consistent with prior studies showing that neutralizing antibodies to ZIKV provide robust protection against ZIKV disease in both immune compromised mouse models and mouse models of ZIKV fetal infection (36,37).

Given the robust protection provided by X2.L1 ZIKV induced neutralizing antibodies, we next evaluated the role of vaccination with X2.L1 ZIKV in the protection from lethal ZIKV challenge. We found that vaccination with X2.L1 provided 100% protection from lethal ZIKV challenge in type I interferon knockout mice. These data provide proof-of-concept vaccine strategies that targeted disruption of terminal xrRNA structures of flaviviruses is an important potential target for development of live-attenuated vaccines for this important group of viruses. Several ZIKV vaccine candidates are in different stages of development using different vaccine platforms including DNA based vaccines (38–42), mRNA vaccines (43), virus-like particles (44,45), live-attenuated vaccine candidates (46), and many others. Due to the possible risk that ZIKV infection can enhance overlapping dengue virus outbreaks (47,48) it is critical that ZIKV vaccine candidates produce long-lasting, high-titer, virus-specific, neutralizing antibodies to prevent potential disease enhancement. We have shown that X2.L1 ZIKV produces high-titer neutralization antibodies after a single vaccination at 21 days. Additional studies are needed to define the benefits of this live-attenuated approach to provide long-lasting, high-titer neutralizing antibodies that reduce future risk of dengue virus disease enhancement in endemic regions.

Some of the potential weaknesses of our proposed attenuation approach include the need to evaluate X2.L1 ZIKV in disease enhancement models for dengue virus. While this would be important to test in cell-culture based assays, it is not clear what these studies would provide for in vivo relevance. Additionally, these studies did not evaluate the duration of neutralizing antibody titer over time. Future studies will need to define the duration and protection of X2.L1 ZIKV over time. Lastly, our studies did not evaluate attenuation, safety, and protection in murine models of fetal infection. These studies were beyond the scope of the initial characterization of X2.L1 ZIKV in our murine models but will be important future directions of this ongoing work to develop novel live-attenuated vaccines for ZIKV.

Overall, this study is the first to use targeted structural mutations of xrRNA2 to attenuate Zika virus. We found that xrRNA2 exhibits increased selection pressure to maintain sfRNA production compared to the xrRNA1 structure. We also found that targeted xrRNA2 structural mutations significantly impaired ZIKV virulence and increased sensitivity to type I interferon. Despite X2.L1 ZIKV attenuation in cell culture and in murine models, the virus produced robust neutralizing antibody response that provided protection from lethal ZIKV challenge in immune compromised murine models of ZIKV disease. These studies provide a template for future studies to determine the individual role of xrRNAs in the pathogenesis of flavivirus pathogenesis in a cell-type specific manner. Additionally, these studies provide important insight into future approaches for live-attenuated vaccine development for this important group of medically important viruses.

## Materials and Methods

### Cell Lines and Viruses

The following cell lines were used in this study: African green monkey kidney cells (Vero E6), human lung epithelial cells (A549), and Aedes albopictus cells (C6/36). All cell lines were sourced from the American Type Culture Collection (ATCC). Vero and C6/36 cells were cultured in minimum essential medium (1X MEM), which was supplemented with 1mM sodium pyruvate, 1X non-essential amino acids (100X, ThermoFisher Scientific), 100U/mL streptomycin, 10mM HEPES, and 10% fetal bovine serum (FBS). A549 cells were cultured in Ham’s F-12K medium (1X F-12K), which was supplemented with 1mM sodium pyruvate, 1X non-essential amino acids (100X, ThermoFisher Scientific), 100U/mL streptomycin, 10mM HEPES, and 10% fetal bovine serum (FBS). Mammalian cells (Vero, A549) were kept at 37℃ with 5% CO2, and insect cells (C6/36) were maintained at 28℃ with 5% CO2. The viruses used in this study are clone derivatives of the ZIKV PRVABC59 (WT-ZIKV) and human isolate of ZIKV Brazil (WT-ZIKV Brazil), Paraiba 01 Brazil 2015 (GenBank: KX280026.1). These derivatives include an unmutated wild-type (WT) ZIKV and a mutated clone with an xrRNA2 mutation described as the X2.L1 ZIKV.

### Plasmids and Generation of the X2.L1 ZIKV

The entire genome of the ZIKV Puerto Rico isolate PRVABC59 exists in a two-plasmid cloning system that utilizes two pACYC177 vector plasmids (49). Plasmid 1, termed p1-ZIKV, contained the region between the 5’ UTR and nt 3498. Plasmid 2, termed p2-ZIKV, contained the region between nt 3109 and the 3’UTR. The QuikChange II XL Site-Directed Mutagenesis Kit (Agilent, Santa Clara, California) was used to introduce the X2.L1 C10518G, U10519A mutation into the p2-ZIKV plasmid using the following primers: ZIKV-xrRNA2F (5’-AGGGGCTCACAGGCTCCATGGCTTCTTCCG-3’), and ZIKV-xrRNA2R(5’-CGGAAGAAGCCATGGAGCCTGTGAGCCCCT-3’). The X2.L1 mutated p2-ZIKV and p1-ZIKV were amplified via rolling circle amplification using the Repli-g mini kit (QIAGEN) and the X2 mutation in p2-ZIKV was confirmed by Sanger sequencing (Eton BioScience, San Diego California).

### Propagation of ZIKV isolates

The X2.L1 mutant p2-ZIKV and unmutated WT p2-ZIKV were separately ligated to p1-ZIKV using 10X T4 DNA Ligase (New England BioLabs). The RNA genomes for the ZIKV X2.L1 mutant and WT ZIKV was transcribed from the ligated plasmid DNA using the HiScribe T^7^ ARCA mRNA kit (New England BioLabs). 40 ug of mRNA was then transfected into Vero cells with the MessengerMAX lipofectamine transfection reagent (Invitrogen). Transfected cells were kept at 37℃ with 5% CO2, until approximately 70-80% of cell clearance was observed. The infectious supernatant was spun down to remove any cellular debris, and the clarified supernatant was supplemented with 20% FBS and 20mM HEPES, followed by immediate storage at -80℃. Transfected Vero stocks of both X2.L1 ZIKV and WT-ZIKV were subsequently passaged into C6/36 cells to obtain a higher titer stock virus that was used for downstream characterization experiments.

### Plaque Forming Unit (PFU) and Focus Forming Unit (FFU) Assays

Vero cells supplemented with complete MEM were plated in a 24-well plate at seeding density and then incubated at 37C for 4-12 hours. At the time of infection, virus stocks were serially diluted in 2% FMS, complete MEM. Media was dumped off cultured vero cells and 200uL of virus inoculum was added to each well. Cells and inoculum were incubated at 37C for 1 hour. Following incubation, 500uL of prewarmed 1.5% CMC in MEM is added to each well directly on top of viral inoculum. Cells were then incubated at 37C and 5% CO2 for 5 days undisturbed. To visualize plaques at the end of the incubation period, overlay was aspirated off and enough 4% PFA to cover cells was added to each well and allowed to incubate at room temperature for 30 mins to 1 hour. After proper duration of time, 4% PFA was dumped off, and wells were washed 3 times with 1X PBS. Wells were stained with 1% crystal violet in 70% EtOH for 1 minute then rinsed gently with water. Plates were then left to dry, and plaques were counted.

Vero cells supplemented with complete MEM were placed in a white bottom 96-well plate at seeding density. Infection was carried out as described above. Cells and inoculum were incubated for 2 days and then fixed as described above. Mouse anti-flavivirus group antigen antibody, clone D1-4G2-4-15(Millipore) was used as a primary antibody and donkey anti-mouse IgG antibody conjugated to horseradish peroxidase (Jackson Research Laboratories) was used as a secondary antibody. TruBlue Substrate (Fisher scientific) was used to stain foci and visualized on C.T.L. immunospot.

Plaques or foci in each well were counted and the viral titer, denoted as plaque-forming units/mL (PFU/mL), was calculated using the following equation:

PFU/mL = (number of plaques) x (dilution factor) / (viral inoculum in mL)

The PFU/mL for each well was averaged with the other wells on the same plate to determine the viral titer for that plate. The overall viral titer was calculated as the average PFU/mL of all plates infected with the virus.

### Viral Genome Copy Quantification

Viral RNA was extracted from both the supernatant and cell lysate of infected cells with the E.Z.N.A. viral RNA kit (Omega Bio-Tek). Extracted RNA samples were then quantified by nanodrop. A normalized amount of RNA (500ng – 1000ng depending on experimental rate limiting sample) was reverse transcribed using iScript cDNA synthesis kit (Bio-Rad). Genome copies were quantified via the following reaction: Luna universal probe qPCR master mix (NEB), synthesized cDNA, primers: Zika 1087: (5’-CCGCTGCCCAACACAAG-3’), Zika 1163c: (5’-CCACTAACGTTCTTTTGCAGACAT-3’), and FAM probe Zika1108FAM: (5’-AGCCTACCTTGACAAGCAGTCAGACACTCAA-3’).

### In Vitro Viral Growth Assays

A549 cells were seeded into 6-well plates at a density of 2 x 10^5^ cells/well 3-12 hours prior to infection. Cells were then infected with either WT-ZIKV or X2.L1 ZIKV at a multiplicity of infection (MOI) of 0.1 for 1 hour at 37℃ with 5% CO2 and then washed once with PBS to remove any non-internalized virus. Each well was then supplemented with Ham’s F-12K medium and kept at 37℃ with 5% CO2 until the appropriate time point. At 0-, 24-, 48-, and 72-hours post infection, supernatant and cellular lysate samples were collected for each virus and used for quantification of viral genome copies via RT-qPCR and for quantification of infectious virus via plaque-forming unit or focus forming unit assay.

### 3’ UTR Sequencing to Confirm Presence of X2.L1 Mutation

Viral genomic RNA was isolated from viral stocks using the OmegaBioTek E.Z.N.A viral RNA kit. Using the linker sequence and reverse transcription primers in the **Table**, viral cDNA was generated following the linker pre-adenylation, linker ligation, and reverse transcription protocols as previously described (50). 3’UTR fragments were amplified using the NEB Phusion Polymerase kit with the following parameters: 98C for 30seconds, 30 cycles of 98C for 10 seconds, 58C for 15 seconds, 72C for 15 seconds, 72C for 10 minutes. Confirmation of sequences were determined via Sanger sequencing (Eton BioScience; San Diego, California).

**Table:**
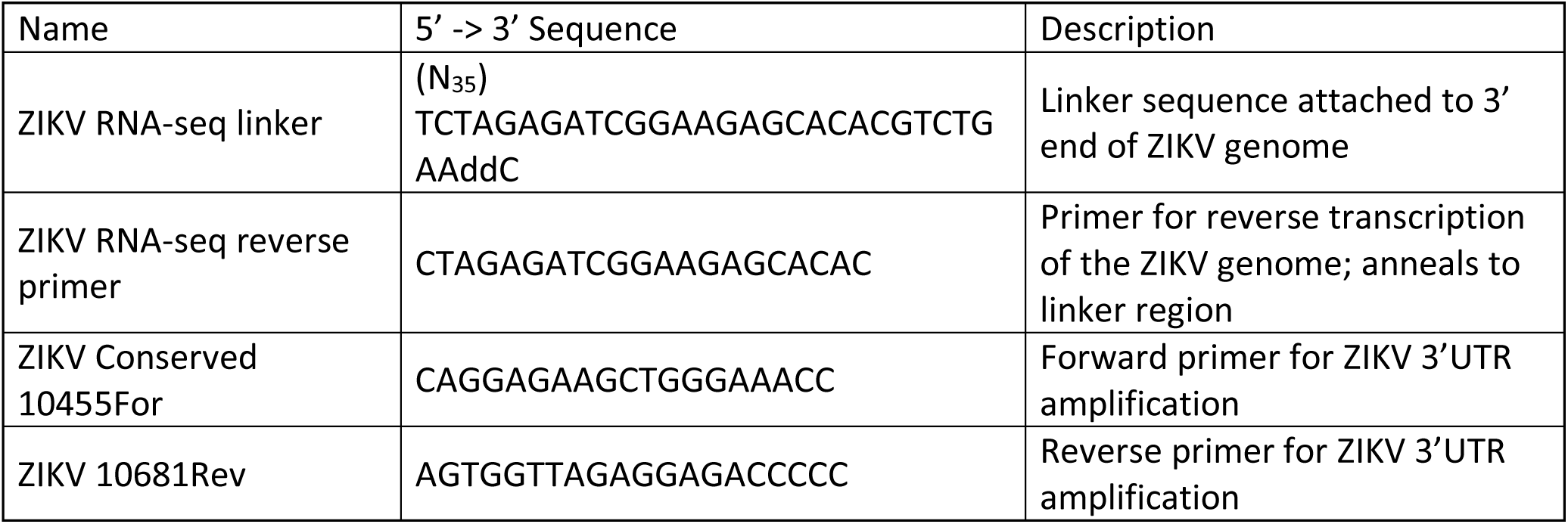
Primer List.

### Interferon Inhibitory Concentration Assays

Vero cells supplemented with complete MEM were plated in a 96-well tissue culture treated plate at 5e^3^ cells per well and left to adhere for 24 hours at 37C. At the time of infection, vero cells were infected at a MOI of 1 in 100uL of inoculum and incubated at 37C for 1 hour. During incubation of virus, interferon-alpha2 (ThermoFisher) was serially diluted 1:2 starting at 200 IU/mL in complete MEM, with the 8^th^ dilution receiving no IFN for a non-IFN treated control. After allotted incubation time, inoculum was removed, and the cells were washed 3 times with PBS. 100 uL of IFN dilutions were then plated directly on to cells and incubated at 37C for 48 hours. After 2 days, 50uL of complete MEM was added to each well to bring final volume up to 150uL so RNA extractions can be performed as previously described.

### Cytotoxicity Assays

Yellow tetrazolium salt (XTT) assay (Invitrogen) was performed as previously described (23). In brief, A549 cells were pated at 10^4^ in a clear bottom, 96-well tissue culture treated plate. Cells were infected at an MOI of 0.1 with virus of interest for 1 hour, then complete MEM was added to a final volume of 200ul/well and incubated until desired time point. At desired time point 50uL of XTT solution was added to each well and incubated at 37C for 4 hours. The plate was then quantified by measuring absorbance at 450nm on VersaMax plate reader.

### 3’ UTR Xrn1 Digest Assay

Wt-ZIKV and X2.L1 ZIKV RNA underwent PCR amplification of the 3’UTR as previously described (23). Using a primer downstream of xrRNA1, a truncated 3’UTR RNA that excluded xrRNA1 was then folded by heating to 65C – 95C for 2-5 minutes then slowly cooled. XRN1 and the folded RNA product where then incubated together for 60 mins at 37C. Following 60 min incubation the product was then run on a 1% agarose gel at 150V for 1.5 hours and visualized on a Biorad Gel Imaging system.

### Animal Studies

All animal and infectious disease studies were reviewed and approved by the UT Southwestern Medical Center Institutional Animal Care and Use Committee and Safety Committee. IFNAR1 KO and AG129 mice were purchased from Jackson laboratories and were maintained and bred in specific pathogen-free facilities at the UT Southwestern Medical Center animal facility. Animals infected with ZIKV were housed in animal BSL-2 facilities. After infection, animals were observed daily for weight changes and signs of disease until meeting endpoint criteria or end of the experimental endpoint. For evaluation of disease-free survival, 6- to 8-week-old IFNAR1-KO mice were infected intraperitoneally (ip) with 10000 pfu of either WT-ZIKV Brazil or X2.L1 ZIKV. Mice were then monitored daily for weight loss and signs of illness including odd behavior, lethargy, neurological impairment, laps in self-hygiene, and any other physical abnormalities. If severe symptoms were observed or mice reached 80% of their original body weight (net 20% weight loss), they were euthanized with isoflurane.

### Murine ZIKV Infection Model

8-week-old IFNAR1-KO mice were infected with 10000 pfu ip with either X2.L1 or PBS (mock). After 21 days mice were infected with 10000 pfu of WT-ZIKV Brazil subcutaneously (sc). Mice were monitored daily checking for signs of illness including odd behavior, lethargy, neurological impairment, laps in self-hygiene, and any other physical abnormalities. They were also weighed daily and euthanized if they reached 80% of their original weight (net 20% weight loss). If mice were observed to have severe symptoms or significant weight loss, they were euthanized with isoflurane.

IFNAR1-KO (Jax #028288) mice or Ag129 (Jax #029098) mice between 6 and 8 weeks of age were infected with 10000 pfu of virus via IP injection. After 8 days mice were euthanized using isoflurane overdose, profused with 30mL of sterile 1X PBS, and spleens and brains were collected and stored in RNAlater (ThermoFisher #AM7021) or 1X PBS depending on further quantification. Organs stored in RNAlater were processed for viral genome quantification via RNA extraction using omega total RNA kit 2, 1000ng of RNA was then processed as previously described. Organs stored in PBS were weight, suspended in 1mL of 1X PBS, manually dissociated with dounce homogenizer, then passed through a 70um filter. Samples were then tittered using the methods previously described.

### Collection of mouse serum

8-week-old IFNAR1-KO mice were inculcated with 10000 pfu/mouse IP of respective virus. After 21 days mice were euthanized as previously described and whole blood was collected via cardiac sick. Blood was then left to coagulate at room temperature for 45 minutes. Once clot is formed samples are spun at 1000-2000 xg at 4 C for 10 minutes. After centrifugation, top serum layer is transferred to a new tube and stored at –80C until use. Serum was pooled from mice inoculated with X2.L1 ZIKV. 100uL of pooled sera or sera collected from uninfected IFNAR1 KO mice was given to ag129 mice IP. After 24 hours mice received 10000 Pfu of WT-ZIKV Brazil subcutaneously. Mice were then monitored for the next 15 days for weight loss and symptom onset.

### Antibody neutralization assay

Neutralization assay preformed as previously described (14). In brief, 10,000 vero cells were plated per well in Nucleon white bottom 96-well plates. Heat inactivated serum collected from infected animals was serially diluted then incubated at 37C for 1 hour with 10 PFU/well WT-ZIKV Brazil. After 1 hour inoculum was plated onto previously prepared Vero’s and incubated at 37C for 1 hour. After 1-hour, warmed CMC was added to all wells and plates incubated at 37C for 40-48 hours then stained as described above.

### ELISA Antibody Studies

Zika Virus Total Antibody Detection Assay Development Kit from Native Antigen company (Cat # ELS61258) was purchased and used to determine total antibody to ZIKV. Serum samples (collection described above) were diluted 1:2, 1:4, 1:8 and 1:16 in duplicate and ELISA was performed as described by the manufacturer protocol.

## Acknowledgements

The authors thank Jeffery S. Kieft, Monica E. Graham, and Elizabeth Spear for their support during early work for this manuscript with northern blot analysis.

## Notes

### Competing Interest Statement

The authors have declared no competing interest.

### Funding

- National Institute of Allergy and Infectious Diseases: R01AI153724
- National Institute of Neurological Disorders and Stroke: R01NS123431

